# Focal cortical surface cooling is a novel and safe method for intraoperative functional brain mapping

**DOI:** 10.1101/2020.05.20.104364

**Authors:** Kenji Ibayashi, Araceli R. Cardenas, Hiroyuki Oya, Hiroto Kawasaki, Christopher K. Kovach, Matthew A. Howard, Michael A. Long, Jeremy D.W. Greenlee

## Abstract

**Objective:** Electrical cortical stimulation (ECS) has been the gold standard for intraoperative functional mapping in neurosurgery, yet it carries the risk of induced seizures. Here we assess the safety of focal cortical cooling (CC) as a potential alternative to ECS for functional brain mapping.

**Methods:** We retrospectively reviewed 40 consecutive subjects (n=13 tumor, 27 mesial temporal lobe epilepsy (MTLE) resection) who underwent intraoperative CC during craniotomy at the University of Iowa Hospital and Clinics from 2007 through 2019 (CC group). Thirty-eight of the 40 subjects had ECS performed along with CC during the same procedure. To assess the safety of CC, intra- and post-operative seizure incidence and post-operative neurological deficits were collected together with new post-operative radiographic findings not related to the surgical procedure itself (i.e. non-mapping portions). As a control cohort, we collected 55 consecutive subjects (n=21 MTLE, 34 tumor/vascular pathology) who underwent awake ECS mapping without CC between 2006 and 2019 (ECS-alone group). To evaluate potential long term effects of mapping techniques (CC and/or ECS), we separately collected another 25 consecutive subjects who underwent anterior temporal lobectomy(ATL) without CC nor ECS between 2007 and 2019 (No ECS/No CC-ATL group).

**Results:** A total of 79 brain sites were cooled in the 40 CC subjects, including inferior frontal gyrus (44%), precentral gyrus (39%), postcentral gyrus (6%), subcentral gyrus (4%) and superior temporal gyrus (6%). No intraoperative seizures were reported in the CC group, whereas 3.6% of ECS-alone group had intraoperative seizures. The incidence of seizure(s) within the first post-operative week did not significantly differ amongst CC (7.9%), ECS-alone (9.0%) and No ECS/No CC-ATL groups (12%). There was no significante difference in the incidence of postoperative radiographic change between CC (7.5%) and ECS-alone groups (5.5 %). The long term seizure outcome for MTLE subjects did not statistically differ regarding ‘good’ outcomes (Engel I+II): CC group (80%), ECS-alone (83.3%) and No ECS/No CC-ATL group (83.3%).

**Conclusions:** Cortical cooling when used as an intraoperative mapping technique is safe, and may complement traditional electrical cortical stimulation.

## INTRODUCTION

Intraoperative functional mapping is a well-established technique in neurosurgery. After introduction in the late 19^th^ century followed by its practical utilization in the 1930s^1^, electrical cortical stimulation (ECS) during awake craniotomy has been the gold standard for reversible cortical perturbation, allowing delineation of functionally-critical cortical sites and guiding surgical resections^1^. ECS has contributed to the understanding of the functional organization and connectivity patterns in the human brain^2^ and provided knowledge to improve the safety and outcome of surgical procedures^2,3^. Despite its great contributions, one drawback of ECS is that large electrical currents are sometimes required to produce observable behavioral changes; these large stimulus intensities increase the risk of intraoperative seizures which can preclude the opportunity to continue the mapping procedure^4,5^. Stimulation-induced seizures can happen even when stimulation intensities are below after-discharge thresholds as detected by concurrent electrocorticography (ECoG)^6,7^. Another limitation with ECS is the difficulty in understanding and estimating the current spread. For example, perturbations in synaptic activity as well as fibers of passage result from ECS and therefore stimulation-induced local and distant network effects are likely^8,9^. This mechanistic uncertainty can complicate the interpretation of stimulation-induced behavioral changes.

Cortical cooling (CC) is a technique introduced and investigated for about a century. CC has proven to be effective in reducing seizure activity ^10-12^ by supressing the neocortex reversibly^13^. Its utility and safety to control cortical excitation has been reported both from experimental animal^11,14,15^ and human research^10,16^. A common form of CC clinical utilization is topical application of cooled saline during events of epileptiform discharges and seizures during neurosurgeries^17-20^. Another method of CC is direct and focal tissue cooling via a surface cooling probe^21,22^. Focal CC in the human brain has been reported previously, noting that the effect is limited within 4mm from the neocortical surface, is reversible, and decreases the power spectrum of the local ECoG^21^.

Together with evidence from animal studies that CC can reversibly alter neural function as does ECS^15^, recent studies have applied CC to investigate human cortical function.^22,23^ Reports show CC can alter behavioral performance in a more graded fashion than ECS^22^. This suggests that CC may perturb cortical function in a manner different than ECS and therefore offer different cortical mapping possibilities that might be able to complement ECS during neurosurgical procedures. Here we assess the safety of CC used for intraoperative cortical mapping during neurosurgery. This is a necessary first step to evaluate CC as a novel complement for ECS; our results show that CC is a safe method to apply during neurosurgical procedures.

## METHODS

### Subjects

This study was approved by the University of Iowa (UI) Institutional Review Board. Subjects were neurosurgical patients who required craniotomy for treatment of either medically intractable epilepsy or intra-axial tumor between 2007 and 2019. There were 56 subjects enrolled in this study and 40 subjects completed CC techniques during surgery (CC group). Within the CC group, 38 subjects (95%) also had intraoperative ECS (ECS/CC group) and 2 subjects (5%) had CC alone. Thirty-one CC subjects (77.5%) underwent awake craniotomy (aCC group) and 9 subjects (22.5%) had craniotomy under general anesthesia (gCC group). Note that 29 of the 31 aCC subjects had ECS performed in the same operation. All of the gCC subjects had ECS performed in the same setting. Of the 40 subjects undergoing CC, 27 (67.5%) underwent anterior temporal lobectomy (ATL) for mesial temporal lobe epilepsy (MTLE), within which 25 subjects had both CC and ECS intraoperatively (ECS/CC-ATL group). The remaining 13 subjects (32.5%) underwent craniotomy for tumor resection in peri-Sylvian areas (Table 1, left column).

**Table1.**
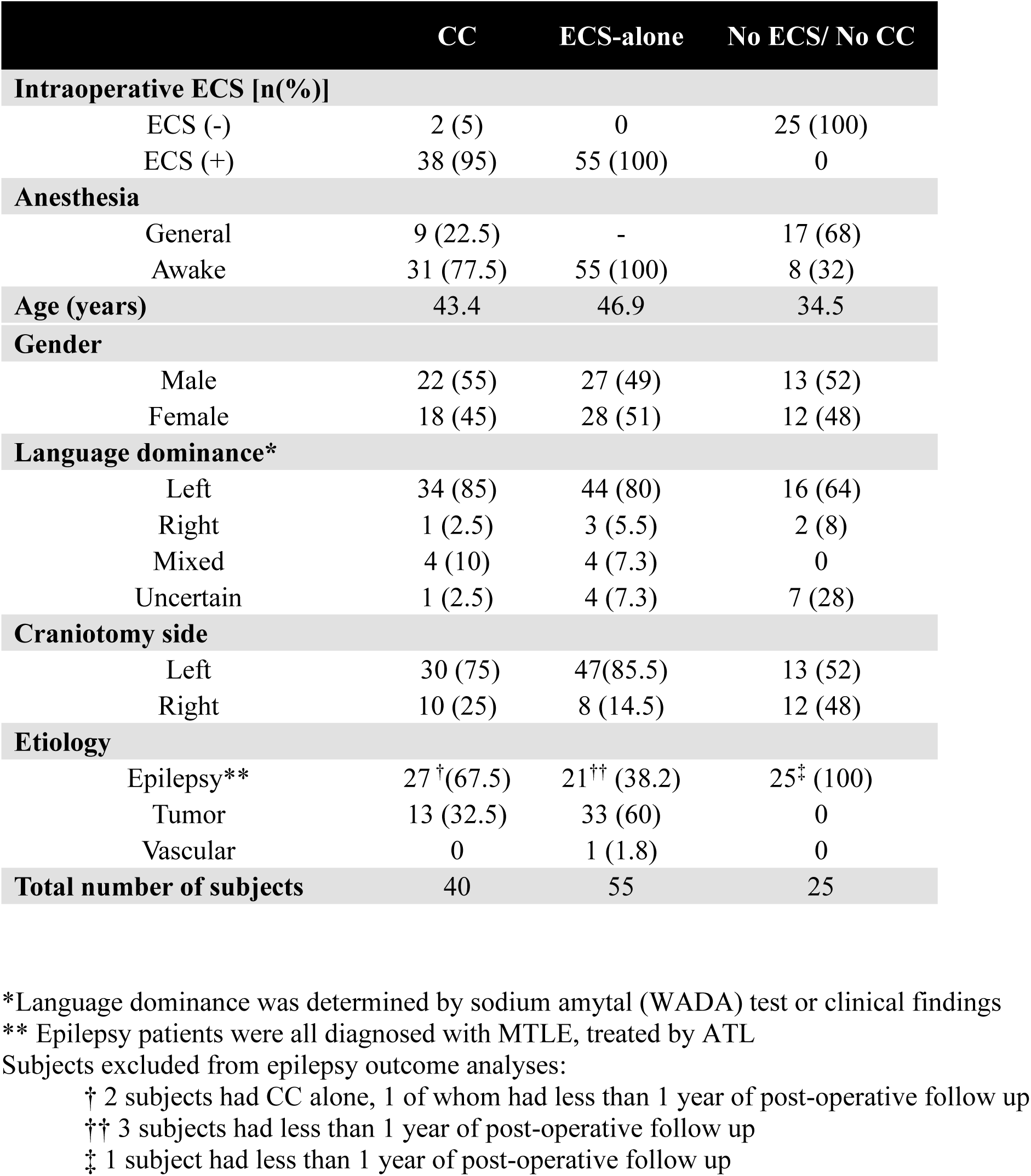
Demographic data

As a control cohort to assess CC safety throughout the perioperative course, we retrospectively collected 55 additional subjects that underwent awake intraoperative ECS mapping without CC (ECS-alone group). This group comprised 21 subjects (38.2%) who had ATL for refractory MTLE (ECS-ATL group) between 2000 and 2019; 18 of these subjects had >1 year follow up. The other 34 control subjects (61.8%) underwent resection surgery for neoplastic or vascular pathologies (33 tumors, 1 AVM) between 2009 and 2019 (Table 1, middle column). Finally, to assess any potential longterm impact of either CC or ECS on the outcome of refractory epilepsy subjects, we retrospectively collected an additional 25 subjects who underwent ATL for MTLE but did not have either ECS or CC during surgery (No ECS/No CC-ATL group). These were subjects treated between 2007 and 2019; one subject was excluded due to less than 1 year follow-up after surgery, so 24 subjects in this second control group were used for subsequent analyses (Table 1, right column).

Age, language dominance, craniotomy side, and etiology are provided in Table 1. Cerebral dominance for language was determined for each subject based on preoperative sodium amytal (WADA) test if completed as part of clinical protocol, otherwise this determination was based on the documentation of the clinical record (e.g. handedness, ictal or post-ictal semiology, intraoperative testing, functional MRI and/or neuropsychological evaluation). Surgical procedures were done awake or under general anesthesia based on clinical need for intraoperative language mapping or intraoperative ECoG recordings to delineate the seizure focus. Because of the preliminary understandings of the role, implementation and interpretation of CC, any transient behavioral alterations resulting from CC were not used to guide surgical decision making. All subjects that underwent ECS and/or CC had appropriate antiepileptic drugs (AED) during surgery either by continuation of their prescribed oral AED, or by intravenous administration of AED during surgery.

### Cortical cooling

CC often was targeted at language-critical cortices before a required corticotomy was made. In the awake subjects, CC was performed either during language-related behavioral tasks or as part of passive (i.e. non-task based) electrical stimulation tract tracing (ESTT)^24,25^ to investigate speech-related areas. These procedures were usually preceded by ECS to determine potential sites for cooling. More specifically, language critical sites as determined by ECS were targeted as sites to test with CC. For the subjects under general anesthesia, CC was performed either together with ESTT, or alone without any preceding tasks for the purposes of baseline measurements and refinement of the CC method. The typical target CC cortical / tissue temperature was below 20°C, which is reported to disrupt local synaptic activity in detailed experimental animal study^26^, and our previous pilot study in human^21^. Localization of CC and ECS sites were determined by intraoperative photographs together with pre- and post-operative imaging studies (MRI or CT) to generate patient-specific brain surface renderings (Fig.1).

**Figure 1:**
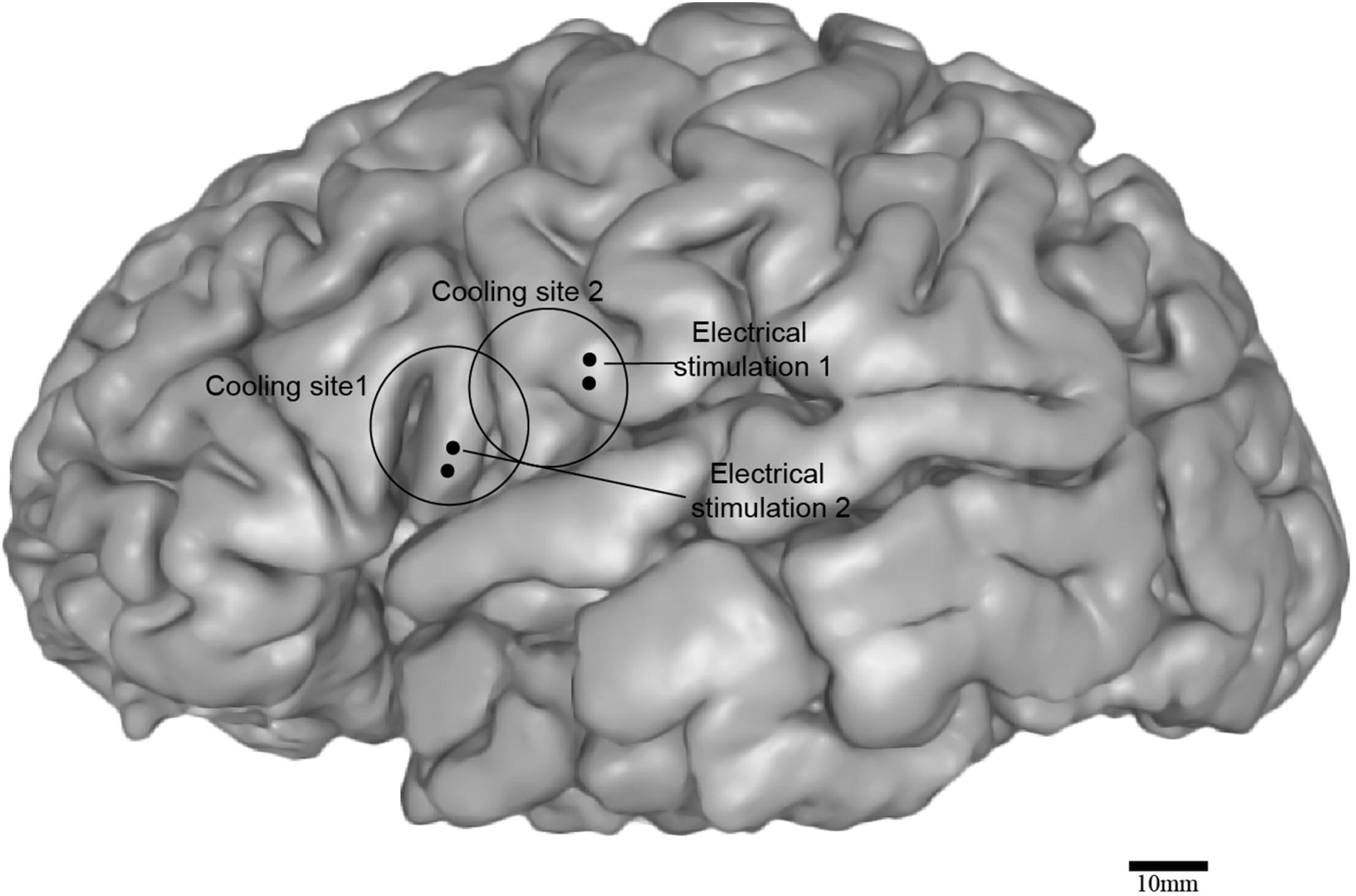
Example of overlapping CC and ECS sites. Lateral surface MRI rendering of an example subject (L187) demonstrating that CC and ECS were often utilized at overlapping / similar cortical regions. This subject had two ECS sites and two CC sites including the precentral gyrus (ECS site 1, CC site 2) and inferior frontal gyrus (ECS site 2, CC site 1).

### Cooling Probes

The cooling devices used in this study have been described previously^22^. In short, two different types of cooling probes were used. One probe was a stainless-steel chamber with a 2 cm diameter circular footprint (Figs.2A,C) and was cooled by infusing sterile chilled hypertonic saline through the chamber. The second probe was a titanium chamber with a 1 × 1 cm square Peltier element (Custom Thermoelectric, Bishopville, MD) attached to the bottom of chamber (Figs.2B,D). Sterile chilled saline infusion through the chamber was used to dissipate heat generated on the non-brain side of the Peltier element. Current to the Peltier element was supplied using a regulated power supply (HY1803D, Tekpower, Taiwan).

**Figure 2:**
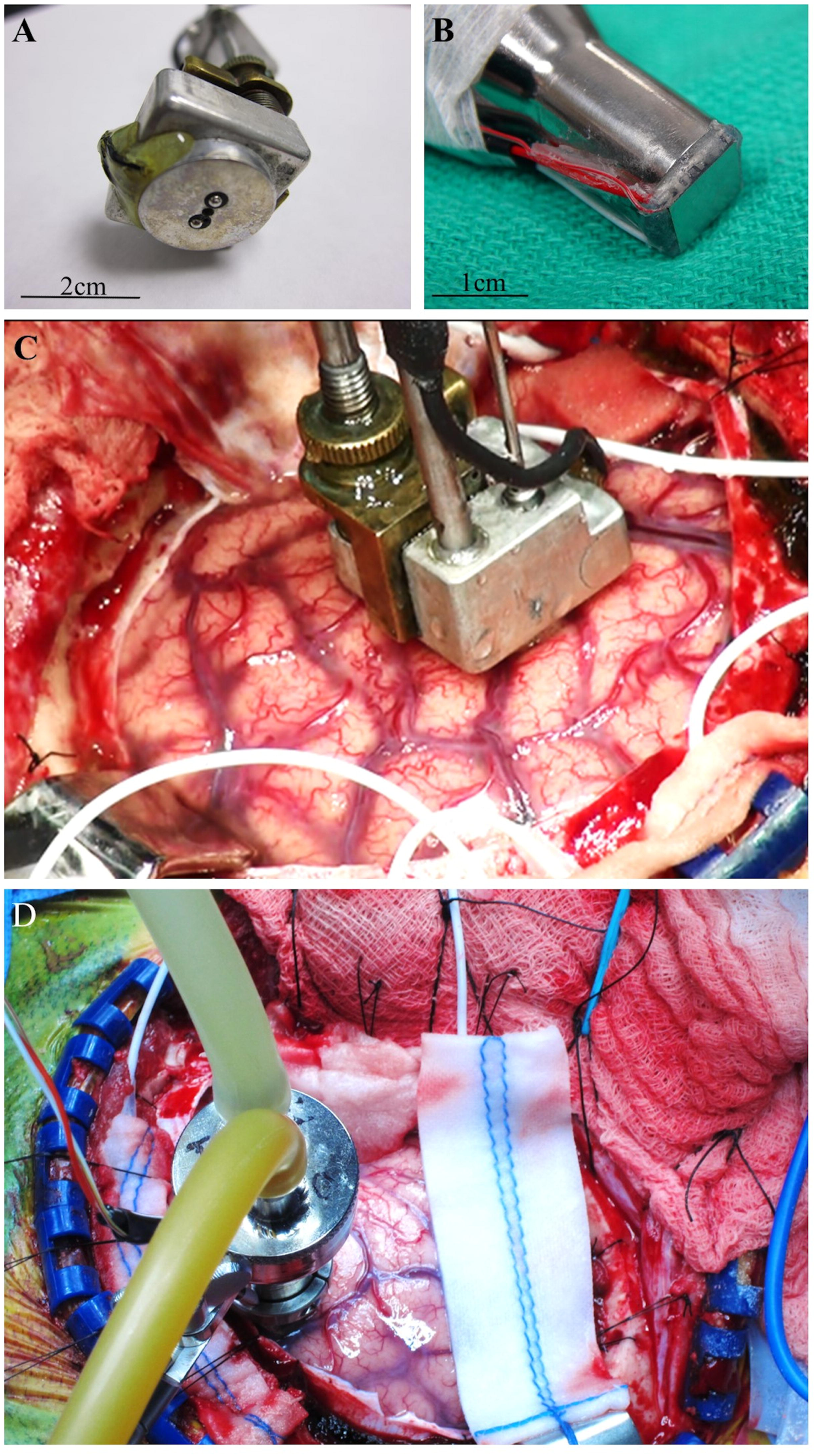
Two types of cooling probes were utilized. Two different cooling probes were used in this study. One probe was a stainless-steel chamber with a 2 cm diameter circular footprint (A, C) which was cooled by infusing sterile chilled hypertonic saline through the chamber. The second probe was a titanium chamber with a 1 x 1 cm square Peltier element (Custom Thermoelectric, Bishopville, MD) attached to the bottom of chamber (B, D), and used sterile chilled saline through the chamber to dissipate heat generated on the non-brain side of the Peltier element.

### Electrical Cortical Stimulation

ECS was performed to identify language-critical cortices during awake surgery using standard clinical techniques and equipment. A custom-made bipolar silver ball-tipped probe was used to deliver 50Hz trains of 0.2ms biphasic, charge-balanced constant-voltage pulses between 5-20V (Grass SD-9, Natus Neurology, Inc., Warwick, RI). Continuous ECoG recordings over cortical areas adjacent to stimulation sites were utilized throughout ECS to monitor for after discharges and determine after discharge thresholds, which defined the upper limit of stimulation intensity. The ECoG recordings were monitored by epilepsy neurologists in the operating room; these physicians were not affiliated with this study.

### Intraoperative evaluation

To evaluate the immediate effects relating to safety during surgery, the incidence of intraoperative seizure events was evaluated retrospectively from the clinical record for all three subject groups: CC, ECS-alone, and No ECS/NoCC-ATL.

### Postoperative evaluation

For delayed safety assessments, post-operative clinical course was evaluated through examinations performed in a pre-discharge inpatient setting as well as longer-term outpatient follow-up for all groups. Subject medical records were retrospectively reviewed. In addition, an IRB-required independent safety monitoring neurology physician not affiliated with this study performed an independent chart review on an annual basis for all subjects undergoing CC. Those safety monitor’s records were reviewed for this report. Finally, perioperative brain imaging studies were reviewed. Appearance of imaging changes (e.g. diffusion restriction) which were otherwise unexplainable related to surgery and lesion resection were evaluated for potential cooling- or stimulation-related complication. Likewise, neuroimaging data from the ECS-alone group was reviewed by the first author to investigate the incidence of post-operative radiologic changes potentially arising from ECS. Any new neurological deficits after surgery were tabulated. The fraction of subjects with positive neurological or radiological findings was compared among the three groups using SPSS (ver.26).

### Post-operative evaluation for seizure outcome in refractory epilepsy patients

To evaluate any impact of CC on long term prognosis for epilepsy patients, we retrospectively collected the seizure outcome based upon the clinical records from the last known visit as of April 2020. Seizure outcomes were based on the Engel classification^27^. The effect of CC on overall seizure control was assessed by comparing ECS/CC, ECS-alone and No ECS/No CC groups for those subjects undergoing ATL and who had at least 1 year follow up following surgical resection. The fraction of ‘good’ outcome (Engel I and II) and seizure freedom (Engel Ia-Ic) was analyzed by Chi-square test of independence implemented in SPSS.

## RESULTS

### Cortical areas that were cooled

Successful CC was accomplished in 40 patients including both aCC and gCC groups. In total 79 cortical sites were cooled; the number of sites ranged from one to seven within subjects (mean = 2.0 ± 1.3 S.D; Fig3; see also Table 2). Among the 79 sites, regions cooled included inferior frontal gyrus (IFG; n=35sites, 44%), precentral gyrus (PrCG; n=31 sites, 39%), postcentral gyrus (PoCG; n=5sites, 6%), subcentral gyrus (SubCG; n=3sites, 4%) and superior temporal gyrus (STG; n=5sites, 6%; Fig3 inset).

**Figure 3:**
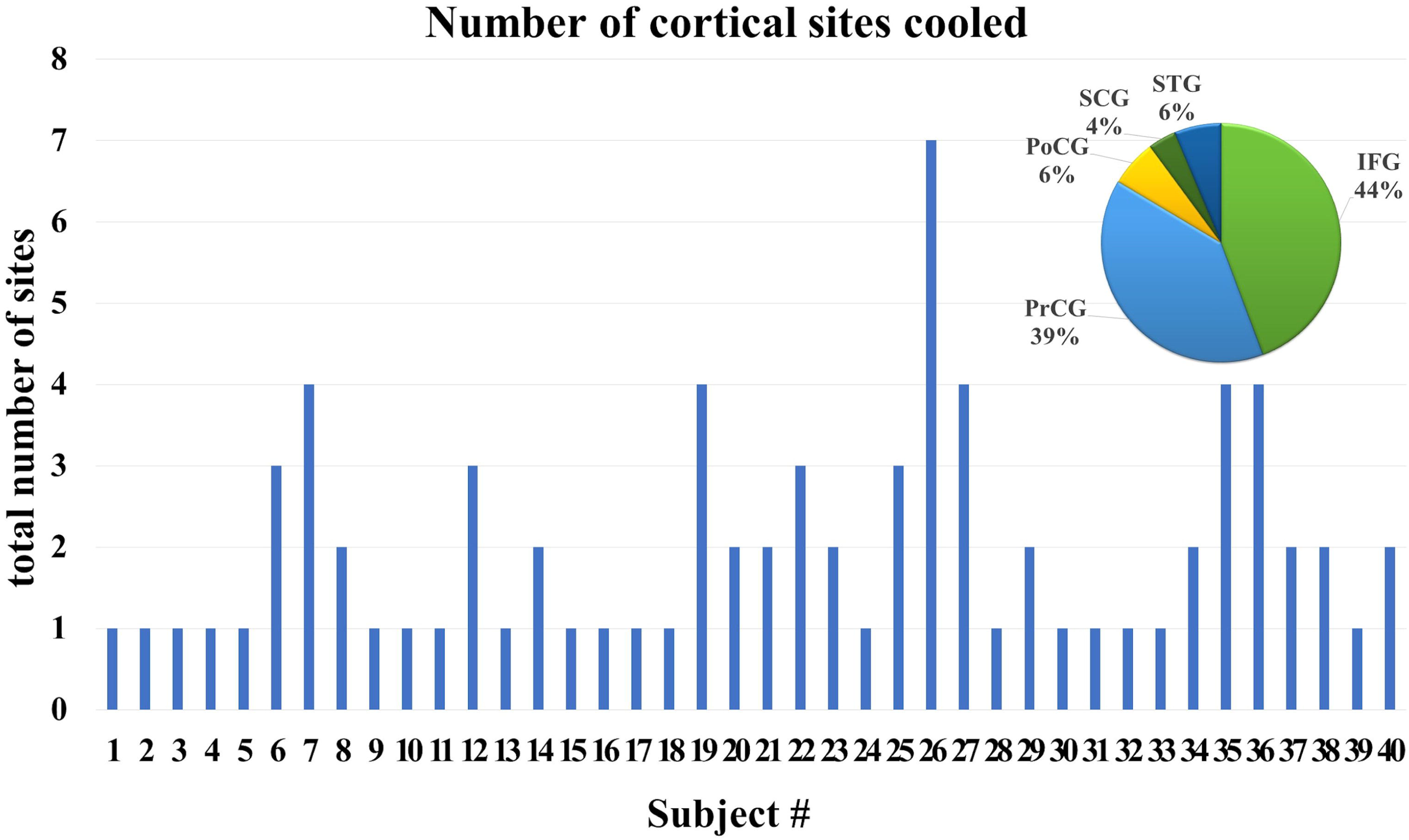
Number and locations of sites cooled. Forty subjects had CC applied during surgery. In this display, subjects 1-9 had CC during general anesthesia (gCC), and the rest had awake craniotomy with CC (aCC). The number of sites cooled per subject varied from 1 to 7 (mean = 2.0 ± 1.3 S.D.). In total 79 cortical sites were cooled including inferior frontal gyrus (IFG; n=35sites, 44%), precentral gyrus (PrCG; n=31 sites, 39%), postcentral gyrus (PoCG; n=5sites, 6%), subcentral gyrus (SubCG; n=3sites, 4%) and superior temporal gyrus (STG; n=5sites, 6%; inset).

**Table2.**
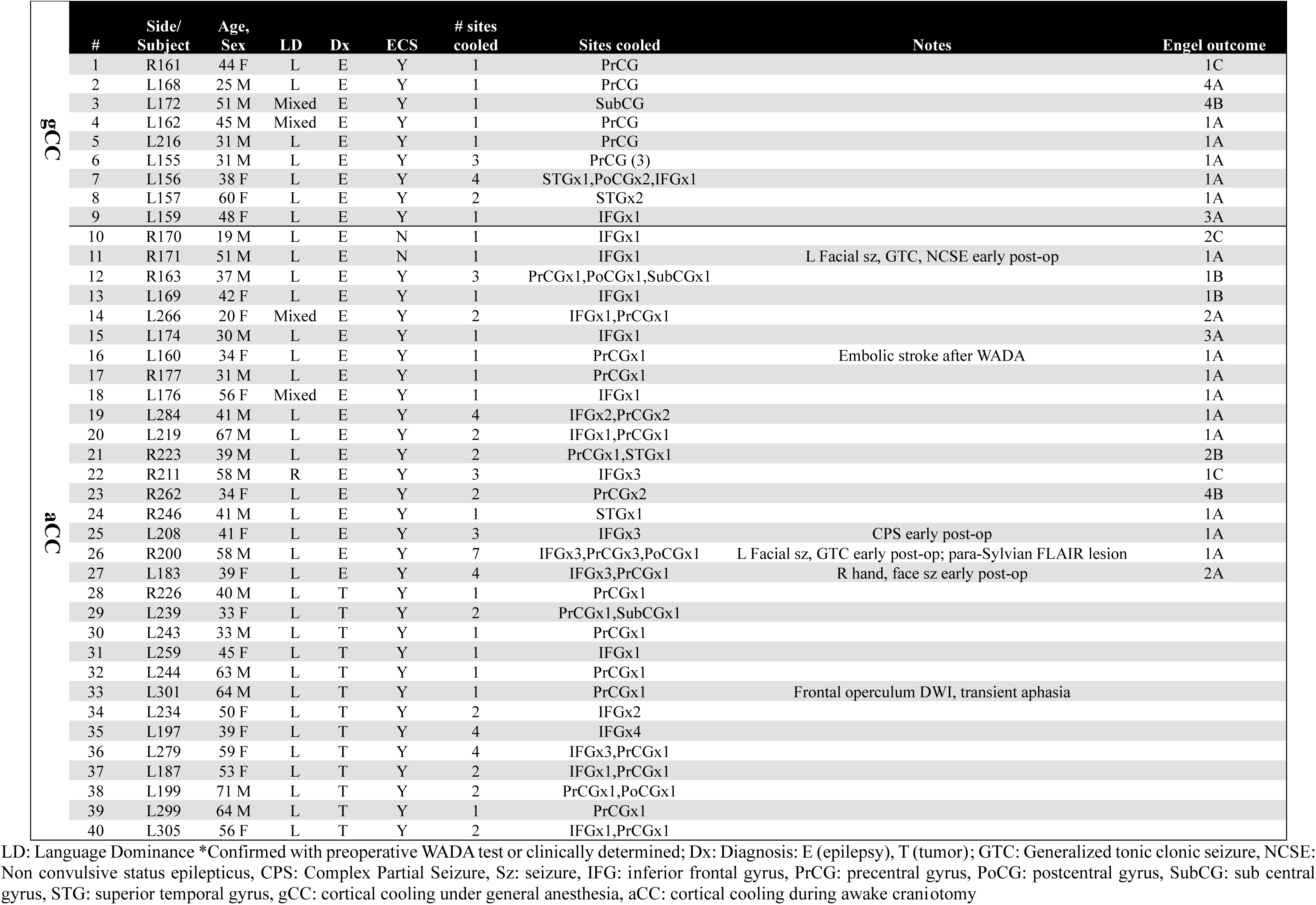
Summary table of all CC subjects

### Intra-operative safety evaluation

Among the 31 aCC subjects, 29 had ECS before CC, of which 26 (90%) had transient behavioral change induced by ECS. The remaining two aCC subjects underwent CC without ECS. There were no intraoperative seizures observed in the CC group. There were 2 subjects (3.6%) who experienced intraoperative seizure in the ECS-alone group. One of these was a subject undergoing left temporal AVM resection who described nausea right after ECS mapping of the temporal surface and cerebral bulging was observed. The nausea was thought to be related to a seizure resulting from the ECS and abated with cold saline irrigation of the brain surface. The other subject was undergoing ATL for epilepsy and manifested habitual seizure during intraoperative ECS. In addition to these, there were two subjects (3.6%) in the ECS-alone group that had asymptomatic prolonged after discharge which ceased with application of cold saline to the exposed cerebrum.

### Post-operative seizures and new neurological deficits

The incidence of within-first-postoperative-week-seizures did not significantly differ between groups: ECS/CC group-7.9%, ECS-alone group-9.0%, and NoECS/NoCC-ATL group-12% (*χ*^2^(2,N=118)=0.17, p=0.92). Furthermore, no differences were identified upon limiting subjects to those undergoing ATL as a more-tailored across group comparison: ECS/CC-ATL cohort (12% post-op seizure incidence; 3 subjects of 25) versus ECS-alone ATL cohort (14.3%; 3 subjects of 21; *χ*^2^(1,N=46)=0.05, p=0.82). Likewise, the incidence of post-operative neurological deficits between ECS/CC and ECS-alone groups was not different (*χ*^2^(1,N=93)=0.09, p=0.76).

Within the first post-operative week, there were four CC subjects (10%; R171, L183, R200, L208) who experienced transient seizures; these ranged from simple facial seizures (L183,R200), complex partial seizure (L208) to nonconvulsive status epiltpticus (R171; Table 2). The four subjects who had seizure episodes were all aCC MTLE subjects and all had ECS in addition to CC except R171 who only had CC. Therefore the incidence of post-operative seizure was 7.9% when limited to ECS/CC group. No gCC subjects had post-operative seizures. There was one aCC subject (L299) treated with metastatic tumor resection who developed episodes of contralateral hand numbness two months after surgery; these were presumed (i.e. no EEG was done) to be seizure by the neurologist and abated with AED. In the ECS-alone group, there were five subjects (9.0%) who developed seizures in the first post-operative week. The semiologies included worsening of expressive aphasia accompanied by continuous electrical seizure activity in the EEG, habitual seizure of right facial numbess and speech difficulty, head convulsion, habitual oral/hand automatism and impaired awareness. In the NoECS/NoCC group, who all underwent ATL, there were 3 subjects (12%) who developed seizures in the first post-operative week. The semiologies included generalized convulsive seizure in one and impaired awareness in two subjects.

Regarding new post-operative neurological deficits, no CC subject experienced a transient or permanent neurological deficit related to CC or ECS at delayed outpatient follow-up (≧4 weeks from surgery). Six subjects (15%) in the CC group did have new neurological deficits but these were considered to be the result of surgical procedure and referable to lesionectomy. These included 3 subjects (7.5%) with memory impairment (tumor resection of left temporal lobe glioblastoma [GBM; L197], left ATL for MTLE [L169, L208]), 1 subject (2.5%) with hemiparesis and dysarthria (tumor resection of left frontoparietal GBM [L244]), 1 subject (2.5%) with fourth nerve palsy (left ATL for MTLE [L159]), and 1 subject (2.5%) with anxiety/depressive symptoms (left ATL for MTLE [L266]). For the ECS-alone group, there were 10 subjects (18.2%) who developed new neurological deficit within the first post-operative week: 6 subjects developed language impairments (4 with glial tumor, 1 with AVM, and 1 with left MTLE), 3 subjects developed motor impairment (1 right facial weakness from left frontal cavernoma resection, 1 right facial and limb weakness from left frontal glioma resection and 1 left side limb weakness due to anterior choroidal infarct following right ATL for MTLE), and 1 subject developed short term memory impairment following left ATL for MTLE. These symptoms were all felt to be related to location of resection and not resultant from ECS.

### Post-operative radiographic findings

The incidence of post-operative radiographic changes did not significantly differ between ECS/CC (7.5%) and ECS-alone groups (5.5%; [*χ*^2^ (1,N=95)=0.16, p=0.69]). All of these patients underwent post-operative MRI (CC: 48% of subjects, ECS-alone: 62%) or CT (CC: 52% of subjects, ECS-alone: 38%) based on standard clinical practice. No subject experienced either a persistent imaging change related to CC or immediate (i.e. <24 hrs) change including post-operative diffusion restriction on diffusion-weighted images (DWI). A single patient (R200, Table 2 #21, Fig 3) had a persistent asymptomatic small FLAIR signal change within the craniotomy and involving both frontal and temporal cortex (Fig 4). This patient underwent a right mesial temporal resection for medically refractory epilepsy and experienced focal facial motor seizures hours after surgery and mapping completion; both CC and ECS were performed. CC was conducted at 7 different sites in suprasylvian cortex only, primarily involving PrCG and IFG (Fig 4A). ECS was conducted at IFG, PrCG, and STG. Another patient (L160, Table2 #16, Fig3) had a low density area in the frontal lobe thought to be an embolic event due to WADA procedure. A third patient (L301, Table2 #33, Fig3) showed diffusion restriction in the ipsilateral frontal opeculum and this was thought to be a result of compromised middle cerebral artery branch flow resulting from the transsylvian surgical prodedure. Similarly, a single subject in the ECS-alone group undergoing resection of a superior posterior frontal anaplastic astrocytoma had a persistent small and asymptomatic FLAIR lesion on the cortical surface away from the resection cavity (Fig 5). There were two other subjects with post-operative radiographic lesions thought to be a result of surgical procedure in ECS-ATL group: ipsilateral temporal base edema consistent with venous infarction after ATL for MTLE, and ipsilateral anterior choroidal infarction after ATL for MTLE.

**Figure 4:**
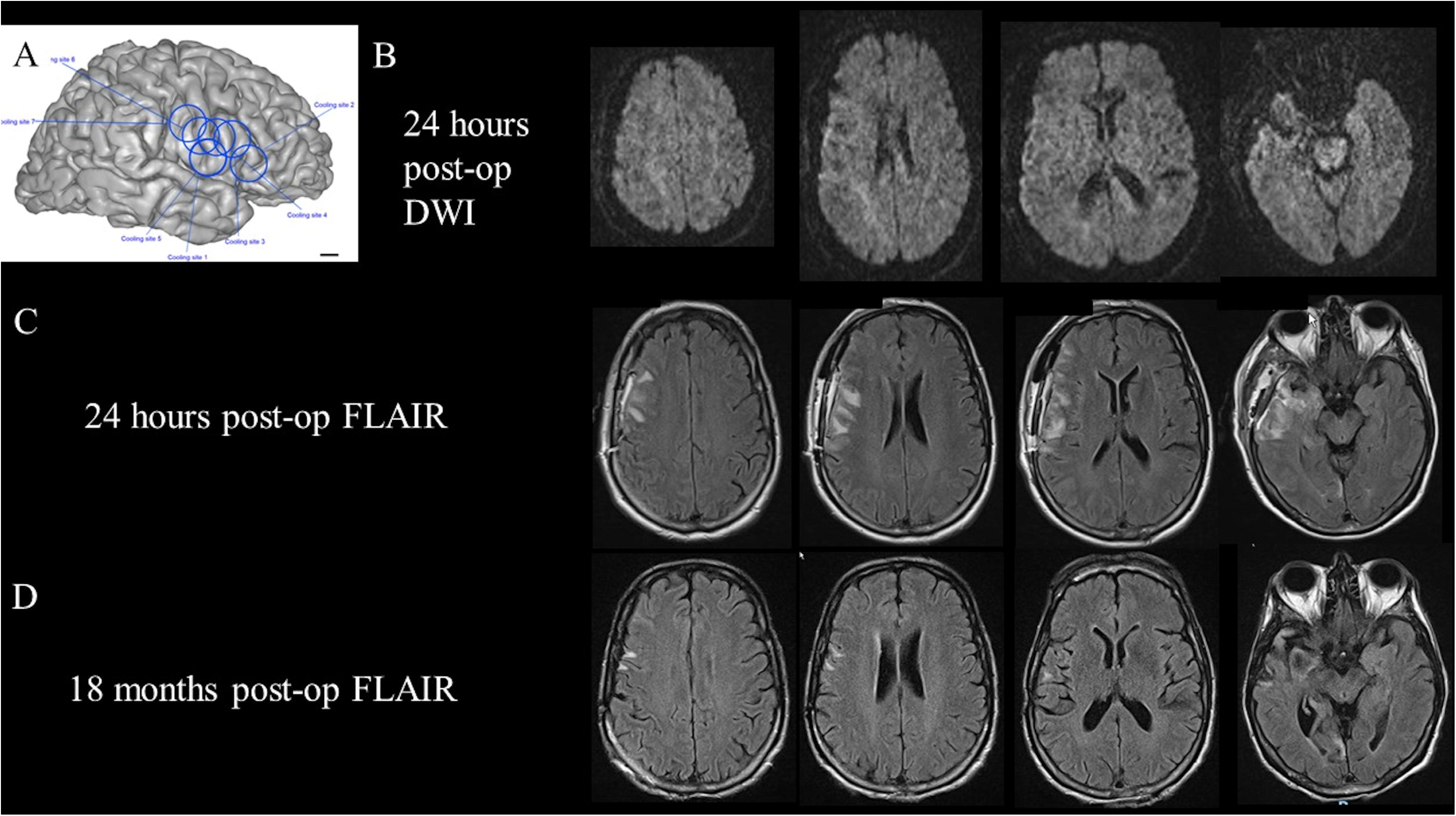
Subject from CC group with MRI changes after surgery. Subject R200 underwent a right mesial temporal epilepsy resection and had CC performed at 7 sites in suprasylvian cortex (A), in addition to extensive ECS mapping. The subject had focal facial seizures and a single generalized seizure which occurred < 24 hrs post-operatively. Early post-op (i.e. <24 hrs) MRI showed no restricted diffusion (row B) but frontal and temporal FLAIR hyperintensities (row C). Delayed imaging 18 months from surgery showed marked improvement in frontal and temporal FLAIR signal change (row D). The patient has remained free of seizures since.POD1 and is neurologically intact.

**Figure 5:**
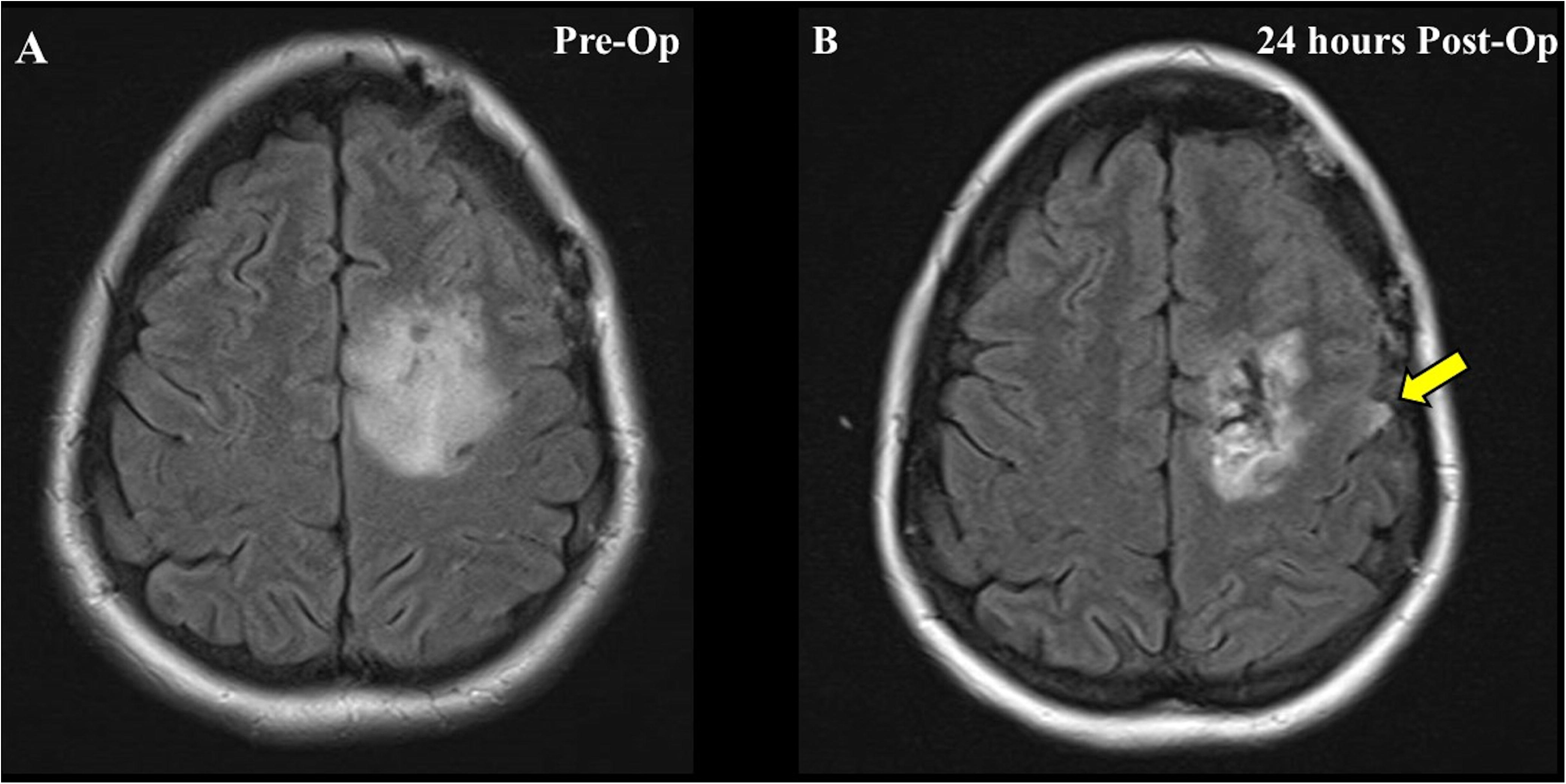
Ssubject from ECS-alone group with MRI change after surgery. This subject underwent left frontal glioma (A) resection with awake mapping procedures. The subject did not have intraoperative seizure, nor prolonged after discharges during the ECS procedure. The early post-operative MRI FLAIR sequence shows a small hyperintense area (B, arrow) on the surface of the ipsilateral precentral gyrus, remote from the resection cavity.

### Post-operative clinical outcome of Epilepsy

The MRI findings, surgical pathology and seizure oucomes are described in Table 3. Note that the CC-alone group had only 1 of 2 subjects with >1 year follow up, so this group was not included in epilepsy outcome analysis. Among the cohorts consisting solely of medically-refractory MTLE who had >1 year post-surgery follow up, the fraction of preoperative abnormal MRI findings in mesial temporal structures (i.e. lesions) were 89% (ECS-alone), 92% (ECS/CC) and 96% (NoECS/NoCC groups) respectively. Similarly, the fraction of positive focal cortical dysplasia in addition to the typical mesial temporal sclerosis were 22% (ECS-alone), 52% (ECS/CC) and 38% (NoECS/NoCC). The ‘good’ outcome rate (Engel I and II) for each group was 83.3% (ECS-alone), 80% (ECS/CC) and 83.3% (NoECS/NoCC) respectively. There was no significant differnce among the three groups (ECS-alone, ECS/CC and NoECS/NoCC [*χ*^2^(2,N=67)=0.12, p=0.94]). The seizure freedom rates (Engel Ia-Ic) were 72% (ECS-alone), 68% (ECS/CC) and 71% (NoECS/NoCC) respectively. There was no significant difference among groups [*χ*^2^(2,N=67)=.10, p=0.95]. The mean follow up duration of the groups were 5.9 ± 4.7, 6.6 ± 3.0, and 4.5 ± 3.6 years (± SD) respectively, and were not statistically different [one-way ANOVA (*F*(2,66)=1.19, p=0.35)].

**Table3.** Impact of intraoperative mapping on the clinical outcome for MTLE after ATL

| Group | Number of Subjects | MRI lesion |  | Pathology |  |  | Engel Outcome |  |  |  | % Good outcome (I+II) † | % Sz freedom (Ia-Ic)‡ | f/u (yrs)†† |
| --- | --- | --- | --- | --- | --- | --- | --- | --- | --- | --- | --- | --- | --- |
|  |  | non lesional | lesional* | negative | MTS | MTS** | I | II | III | IV |  |  |  |
| ECS-alone | 18 | 2 | 16 | 3¶ | 11 | 4 | 15 | 0 | 3 | 0 | 83.3 | 72.2 | 5.9±4.7 |
| ECS/CC | 25 | 2 | 23 | 0 | 12 | 13 | 17 | 3 | 2 | 3 | 80.0 | 68.0 | 6.6±3.0 |
| No ECS/No CC | 24 | 1 | 23 | 0 | 15 | 9 | 17 | 3 | 2 | 2 | 83.3 | 70.8 | 4.5±3.6 |

MTS: Mesial Temporal Sclerosis, Sz: seizure
\* positive preoperative MRI abnormality in mesial temporal structures including amygdala or hippocampus
\*\* positive focal cortical dysplasia outside of MTS
† No significant difference among ECS alone vs CC&ECS vs NoCC/NoECS, $X^2(2, N=67)=0.12$ , $p=0.94$ .
‡ No significant difference among ECS alone vs CC&ECS vs NoCC/NoECS, $X^2(2, N=67)=0.10$ , $p=0.95$ .
†† No significant difference among ECS alone vs CC&ECS vs NoCC/NoECS, one-way-ANOVA ( $F(2,66)=1.19$ , $p=0.35$ )
¶ There were 2 subjects without available pathology reports

## DISCUSSION

This report shares our results with focal brain surface cooling and represents more than 10 years and 40 patients of experience. Our data shows that CC is a safe technique that can be done in both awake and general anesthetic craniotomy settings. While safety of the technique is the focus of this report (and not the ability of CC to induce transient behavioral effects or to guide surgical decision making), we have previously shown CC to be a useful tool to probe eloquent cortex and behavioral function^22^. It is also noteworthy that only two of our subjects underwent CC alone and the remaining 38 of 40 had CC performed along with ECS; therefore, our data cannot purely inform the safety of CC in isolation. To overcome this confound, we included control goups of both ECS-alone as well as patients without any intraoperative mapping that underwent similar surgical procedures to provide context for safety assessments of CC.

### Incidence of Intraoperative Seizures

The current gold standard of any surgical mapping during craniotomy requires perturbation of cortical function to understand the cortex in question’s function. By definition, this alteration will result in changes in cortical excitability and as a result seizure may result. Seizure is one of the well-documented intraoperative complications of ECS, and its prevalence is reported to be between 2.2%^4^ and 21.5%^28^. Seizures have been reported to correlate significantly with the occurrence of perioperative complications such as early and late neurological deficit, hemorrhage, stroke, infection and deep venous thrombosis etc. ^29^. In our study, there were no intraoperative seizures observed in CC group, but two (3.6%) occurred in the ECS-alone group. Animal study of cortical cooling suggests CC to have the potential to induce increased excitability that could generate spikes, but not until the local temperature decreases below 10°C.^26^ While our CC group did not manifest any intraoperative seizure events, this may be due to preventive effect of adequate AED administered before the procedure. In addition, the overall low incidence of intraoperative seizure (including ECS-alone) could reflect a small cohort size, retrospective nature of the study with potential lack of documentation of intra- or peri-operative seizures in the medical record, or could reflect the advancement of neuro-anesthesia or ECoG monitoring techniques.

### Incidence of Post-operative Seizures

We did not identify difference in the incidence of seizures occurring within the first post-operative week for patients undergoing CC compared to our control groups. This result remained even after limiting groups to subjects undergoing ATL in order to optimize the group comparisons. This suggests that CC does not add risk of early post-operative seizure incidence when used together with ECS.

### Post-operative radiological and neurological findings in the CC group

We had one subject in the ECS/CC group who retained a small and asymptomatic residual FLAIR change on longer-term follow up MRI. The fact that the location of this signal change extended beyond the areas actually cooled combined with the known focal (i.e. millimetric) nature of temperature changes we have measured underneath the cooling probe in a laboratory model^22^, suggests it is unlikely that the FLAIR change resulted from CC alone (Fig4C,D), but rather related to the surgical procedure itself or to the post-operative seizures. This patient did experience early post-operative focal and generalized seizures, but did not suffer subsequent longer-term or ongoing seizures. Seizures are known to be a cause of MRI signal change in the post-ictal period.^30,31^. Although we cannot completely exclude the possibility of the culprit being one of or the combination of intraoperative mapping procedures (CC or ECS), the fact that no CC involved the temporal lobe suggests the MRI finding was not resultant from the CC.

We also had one subject in the ECS-alone cohort who developed a small asymptomatic FLAIR signal change adjacent to the surgical field and stimulated cortex (Fig 5). This subject did not have any persistent after discharges or perioperative seizures recorded. The exact etiology of this small MRI finding is uncertain. The incidence of post-ECS MRI changes in human has been under-evaluated in the literature, but there is a report of stimulation-related MRI signal changes in animal kindling models^32^.

No CC subjects exhibited any new neurological deficits after surgery that were attributable to CC. Likewise, there were no permanent deficits related to ECS in both ECS/CC and ECS-alone groups. All the newly developed post-operative neurological deficits were considered to be the result of neurosurgical resection in both ECS/CC and ECS-alone groups based on neuroanatomical, imaging, and clinical correlation. Taken together with the collective peri-operative seizure and post-operative imaging findings, our results support that CC used as an intraoperative mapping technique does not increase the risks to subjects.

### Post-operative seizure outcome for refractory epilepsy patients in CC group

Our results show that CC did not have any negative impact on long term seizure outcome for MTLE subjects. The overall seizure freedom rate of ∼70% matches the previously reported randomized controlled trials and prospective studies for MTLE^33-36^. The current data supports that both CC and ECS, even when early post-operative seizures occur, do not add clinical risk of sub-standard longterm seizure outcomes.

### Limitations and future directions

This retrospective study was not designed to directly compare the efficacy between CC and ECS as intraoperative brain mapping techniques. Additionally, most CC subjects had ECS performed as well during surgery, so our safety data is not a ‘pure’ assessment of CC. Many of the same cortical regions where CC was applied were the same sites where ECS was delivered, and indeed ECS was a faster and more surgically-efficient way to identify behaviorally-relevant sites than CC. To mitigate this confound of combined ECS and CC from a safety standpoint we included a contemporaneous and consecutive series of ECS-alone subjects, as well as subjects who underwent ATL without either of these techniques. Our results support the non-inferiority safety profiles of both ECS/CC and ECS-alone.

Some disadvantages of CC are that it requires longer time and additional equipment in the operative setting, compared to ECS. CC cannot yet target deep structures in the brain. The mechanical modification of the probe to enable deep structure cooling (e.g. a thinner probe for intrafissural cortices or a smaller-sized-probe for transventricular cooling of the hippocampus) could be useful to expand opportunities to map these areas in future. The ability to access deep structures would directly benefit areas of research in relation to seizure activity suppression, given that the mesial temporal structures are a common focus of medically intractable epilepsy. Nevertheless an overall advantage of CC is the capability to acquire a more precise spatial localization of function and the capability to manipulate the behavior while leaving the structure intact instead of eliminating it^15,37,38^, which can lead to greater insight concerning the link between brain function and behavior.

## CONCLUSION

Cortical cooling is a safe and useful technique for intraoperative mapping, and can be used complementarily with electrical cortical stimulation.

## ACKNOWLEDGMENTS

We thank our patients for their participation. We thank Dr. Mark Granner for serving as the independent safety monitor.

